# Centrosome amplification mediates small extracellular vesicles secretion via lysosome disruption

**DOI:** 10.1101/2020.08.19.257162

**Authors:** Sophie D. Adams, Judit Csere, Gisela D’angelo, Edward P. Carter, Teresa Arnandis, Martin Dodel, Hemant Kocher, Richard Grose, Graça Raposo, Faraz Mardakheh, Susana A. Godinho

**Affiliations:** Centre for Cancer Cell and Molecular Biology, Barts Cancer Institute, Queen Mary University of London, Charterhouse Square, London EC1M 6BQ, UK; Structure and Membrane Compartments, Institute Curie, Paris Sciences & Lettres Research University, Centre for National de la Recherche Scientifique, UMR144, Paris, France; Centre for Tumour Biology, Barts Cancer Institute, Queen Mary University of London, Charterhouse Square, London EC1M 6BQ, UK; Department of Pathology, School of Medicine and Dentistry, Catholic University of Valencia, 46001, Valencia, Spain

**Keywords:** Centrosome amplification, exosomes, extracellular vesicles, ROS, lysosomes, PDAC, stellate cells, invasion

## Abstract

Bidirectional communication between cells and their surrounding environment is critical in both normal and pathological settings. Extracellular vesicles (EVs), which facilitate the horizontal transfer of molecules between cells, are recognized as an important constituent of cell-cell communication. In cancer, alterations in EV secretion contribute to the growth and metastasis of tumor cells. However, the mechanisms underlying these changes remain largely unknown. Here, we show that centrosome amplification is associated with and sufficient to promote small extracellular vesicle (_S_EV) secretion in pancreatic cancer cells. This is a direct result due of lysosomal dysfunction, caused by increased reactive oxygen species (ROS) downstream of extra centrosomes. Defects in lysosome function promotes multivesicular body fusion with the plasma membrane, thereby enhancing _S_EV secretion. Furthermore, we find that _S_EVs secreted in response to amplified centrosomes are functionally distinct and activate pancreatic stellate cells (PSCs). These activated PSCs promote the invasion of pancreatic cancer cells in heterotypic 3-D cultures. We propose that _S_EVs secreted by cancer cells with amplified centrosomes influence the bidirectional communication between the tumor cells and the surrounding stroma to promote malignancy.

## Introduction

A variety of human cancer types often exhibit defects in the structure and number of centrosomes, the main microtubule organizing centers in animal cells [1, 2]. Work in fly and mouse models has shown that centrosome abnormalities, in particular centrosome amplification, are not mere byproducts of tumorigenesis but rather play direct roles in promoting and accelerating tumor progression [3–6]. While the full extent by which centrosome abnormalities promote tumorigenesis is still unclear, centrosome amplification can directly promote aneuploidy and cell invasion, which play important roles in malignant progression [7–9]. Recently, we reported that centrosome amplification induces the secretion of several proteins with pro-invasive properties, e.g. interleukin-8, which induces invasive behavior in neighboring cells [10]. This altered secretion is partially due to a stress response that results from increased ROS downstream of centrosome amplification [10]. Thus, the presence of amplified centrosomes can also influence tumors in a non-cell autonomous manner, via protein secretion, suggesting a broader and more complex role for these abnormalities in cancer.

Secretion of cytokines, growth factors and extracellular vesicles (EVs) promote bidirectional communication between cancer cells and the tumor microenvironment (TME). This cross-talk impacts tumor initiation, progression and patient prognosis [11, 12]. EVs are membrane-bound vesicles containing proteins, lipids, DNA and RNA species (microRNA, mRNA and long non-coding RNAs) that can mediate the horizontal transfer of molecules between cells [13]. Their role in cell-cell communication is of particularly interesting due to their long-lasting effects and ability to influence distant tissues, e.g. during pre-metastatic niche formation [14]. Eukaryotic cells secrete two main types of EVs, microvesicles and exosomes, which differ in their size and biogenesis pathways. Microvesicles (large EVs, _L_EVs, ^~^100-1000 nm diameter) are formed through outward budding or “shedding” of the plasma membrane. In comparison, exosomes (small EVs, _S_EVs, ^~^30-150 nm diameter) are generated intracellularly as intraluminal vesicles (ILVs) within multivesicular bodies (MVBs), which are released upon the fusion of the MVBs with the plasma membrane [13]. Both types of EVs are secreted by cancer cells and have been shown to play key roles in tumor progression, potentially via changes in their composition [15, 16].

Exosomes, a subtype of _S_EVs, are critical in shaping the TME [16]. This is particularly clear in the stromal compartment, where cancer-derived exosomes can activate fibroblasts through transfer of molecules such as TGF-β [16–19]. Fibroblast activation leads to the deposition of extracellular matrix (ECM), tumor fibrosis and metastasis [20]. This is particularly important in pancreatic cancer, where activation of the myofibroblast-like stellate cells, and consequent fibrosis, are the major contributors to the highly aggressive nature of these tumors and poor treatment efficacy [21–23]. While some exosomal components are known to contribute to fibroblast activation and recruitment (e.g. TGF-β and Lin28B) [19, 24], the pathways responsible for alterations in their packaging and secretion in cancer cells remain largely unknown.

Here, we show that the presence of extra centrosomes is sufficient to increase secretion of _S_EVs, but not large _L_EVs. Characterization of these _S_EVs by immunoelectron microscopy (IEM) and SILAC proteomic analyses suggests that they are of endocytic origin and thus enriched for exosomes. Mechanistically, we found that disruption of lysosome function, as a consequence of increased ROS in cells with extra centrosomes, prevents efficient lysosome and MVB fusion, leading to _S_EV secretion. Furthermore, _S_EVs secreted by cancer cells with extra centrosomes are functionally distinct and can induce PSC activation. Consequently, pancreatic stellate cells (PSCs) pre-treated with _S_EVs from cancer cells with extra centrosomes promote invasion of pancreatic ductal adenocarcinoma (PDAC) cells in heterotypic 3-D cultures. Our findings demonstrate that centrosome amplification promotes quantitative and qualitative changes in secreted _S_EVs that could influence communication between the tumor and the associated stroma to promote malignancy.

## Results

### Centrosome amplification induces secretion of sEVs

Our previous work demonstrated that centrosome amplification leads to proteomic changes in the secretome, including an increase in proteins associated with EVs, suggesting higher EV secretion in cells with amplified centrosomes [10]. To explore this further, we used an established ultracentrifugation (UC) method [14] to crudely separate EVs according to their size: _L_EVs and _S_EVs, which we validated by nanoparticle tracking analyses (Figures S1A and S1B). To accurately measure secreted EV numbers, we used ImageStream flow cytometry to quantify fluorescently labelled EVs with the lipid dye BODIPY maleimide [25] and ensured that all serum was depleted for existing EVs by UC (Figures S1C and S1D). We found that in the mammary epithelial cell line MCF10A where the secretome analysis was previously performed [10], induction of centrosome amplification, by transient overexpression of the Polo-like kinase 4 (PLK4) in response to doxycycline (DOX) [26], led to increased secretion of _S_EVs, but not _L_EVs (Figure S1E).

Due to the well-established role of _S_EVs in activating fibroblasts, and its downstream effects on pancreatic cancer prognosis and treatment [16, 22], we decided to investigate if the presence of extra centrosomes would impact _S_EVs secretion in pancreatic cancer. To do this, we quantified the number of EVs and percentage of centrosome amplification in a panel of PDAC cell lines. We observed that cell lines with higher levels of centrosome amplification secreted increased numbers of EVs, in particular _S_EVs, demonstrating a significant correlation between extra centrosomes and _S_EV secretion (Figures 1A-1C and S1F). Furthermore, we confirmed that induction of centrosome amplification in two pancreatic cell lines, PaTu-S and HPAF-II, is sufficient to increase secretion of _S_EVs, but not _L_EVs (Figures 1D and S1G). Additionally, depletion of SAS-6, a protein important for centrosome duplication, in cells exposed to DOX and PLK4 overexpression prevented both centrosome amplification and increased _S_EV secretion, suggesting that _S_EV secretion is indeed a consequence of centrosomal alterations (Figures 1D and S1G).

**Figure 1.**
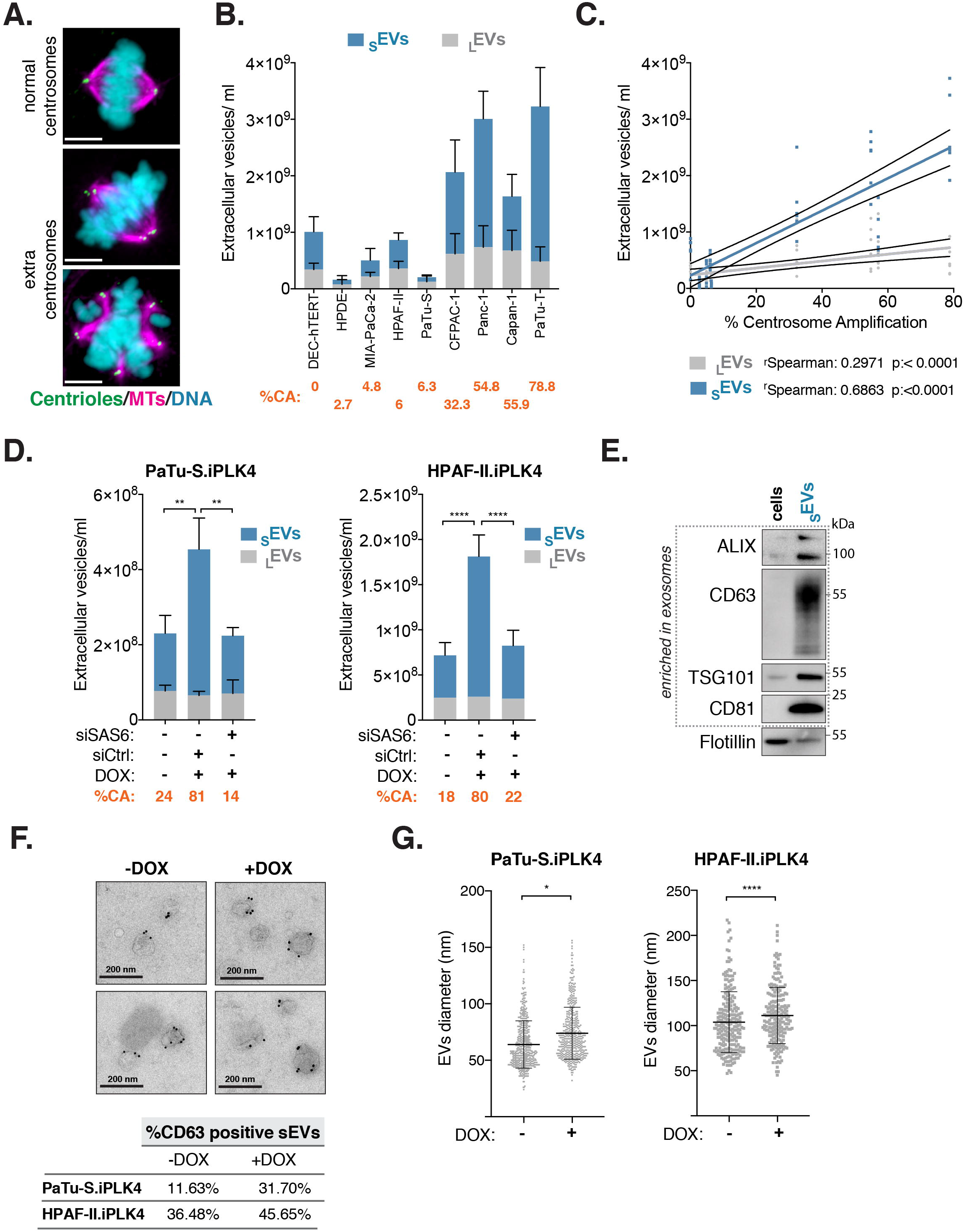
Centrosome amplification promotes the secretion of _S_EVs in PADC cells. (A) Representative confocal images of mitotic cells with normal and amplified centrosomes. Cells were stained for α-tubulin (magenta), centrin2 (green) and DNA (cyan). Scale bar, 10 μm. (B) Quantification of _S_EVs and _L_EVs secreted by PDAC cell lines. Average of the percentage of centrosome amplification (CA) per cell line is highlighted in orange. (C) Linear regression of the data presented in B and Spearman correlation coefficients for _S_EVs and _L_EVs. (D) Quantification of secreted _S_EVs and _L_EVs in Patu-S.iPLK4 and HPAF-II.iPLK4 cell lines upon induction of centrosome amplification (+DOX), before and after depletion of Sas-6 by siRNA. Average percentage of centrosome amplification (CA) per condition is highlighted in orange. (E) Western blot analyses of proteins associated with _S_EVs in extracts from cells and _S_EVs collected by UC. (F) Top: Representative images of IEM of _S_EVs collected from HPAF.iPLK4 cells. Dark beads represent immunogold labelling with anti-CD63. Scale bar, 200 nm. Bottom: Quantification of the percentage of positive CD63 _S_EVs. (G) Quantification of _S_EVs diameter by cry-EM. Patu-S.iPLK4 _S_EVs n_(−DOX)_=232 and n_(+DOX)_=216; HPAF-II.iPLK4 n_(−DOX)_=541 and n_(+DOX)_=493. For all graphics error bars represent mean +/− SD from three independent experiments. \**p* < *0.05,* \*\**p* < *0.01*, \*\*\*\**p* < *0.0001.* The following statistic were applied: for graphs in D two-way ANOVA with Tukey’s post hoc test was applied and for graphs in G unpaired *t* test was applied. See also Figure S1.

The SEVs fractions isolated by UC were enriched for several markers associated with exosomes, such as ALIX, CD63, TSG101 and CD81 [27], but not for general membrane markers, such as flotillin (Figure 1E). We further confirmed the presence of bona fide EVs in the _S_EVs fractions by EM and immunogold labeling using the SEV marker CD63 [28]. Consistent with increased _S_EV secretion, we found that the percentage of CD63^+ve^ EVs was higher in cells with extra centrosomes (+DOX) (Figure 1F). Moreover, these _S_EVs were slightly larger, assessed by EM and also nanoparticle tracking analyses, suggesting that qualitative changes might also occur in these EVs (Figures 1G and S1H). Altogether, our results demonstrate that centrosome amplification promotes _S_EV secretion.

### Proteomic analyses of _S_EVs demonstrates their endocytic origin

To further understand the origin and composition of these _S_EVs, we performed stable isotope labelling by amino acids in cell culture (SILAC) proteomic analyses [29]. SILAC labelling with medium and heavy isotopes enables the exclusion of contaminant serum proteins, which would be unlabeled (equivalent of light labeling), and allows for simultaneous processing of purification steps to decrease sample-to-sample variability (Figure 2A). Because UC isolated fractions can contain contaminants, such as protein aggregates and cellular debris, we further purified the _S_EVs UC fraction using size exclusion chromatography (SEC) prior to proteomics analysis (Figure S2A). Commercially available qEV SEC columns designed to purify exosomes were used [30, 31] and _S_EVs were quantified by ImageStream, as before. As expected for these columns, _S_EVs collected from PaTu-S.iPLK4 cells (−/+ extra centrosomes) eluted in fractions 7-10, with the majority eluting in fractions 8 and 9 (Figure S2B). SILAC reverse and forward labelling was performed to conduct proteomic analyses of fractions 7, 8 and 9. Quantitative analyses of the proteomic data for each SEC fraction revealed that approximately 464 proteins were common to all fractions, and included known _S_EVs components such as ALIX, TSG101, CD81 and CD9 (Table S1). There were also proteins unique to each fraction suggesting that these _S_EVs are heterogeneous (Figure 2B). Comparison of our _S_EV proteomics data with the EV database Vesiclepedia [32] revealed that the majority of proteins in our datasets have been previously identified in other EV studies, confirming the robustness of our purification protocol. Enrichment analyses of common proteins present in both SILAC forward and reverse labeling experiments were performed to identify common pathways (Tables S2 and S3). Importantly, the most significantly enriched categories were associated with EV, _S_EV and linked to pathways unique for exosome biogenesis, such as recycling endosome and endocytic vesicles (Figure 2D). Moreover, pathways linked to cell communication, response to stress, pancreatic secretion and immune response were also enriched in our dataset (Figure 2D), indicating that these _S_EV might have diverse functions.

**Figure 2.**
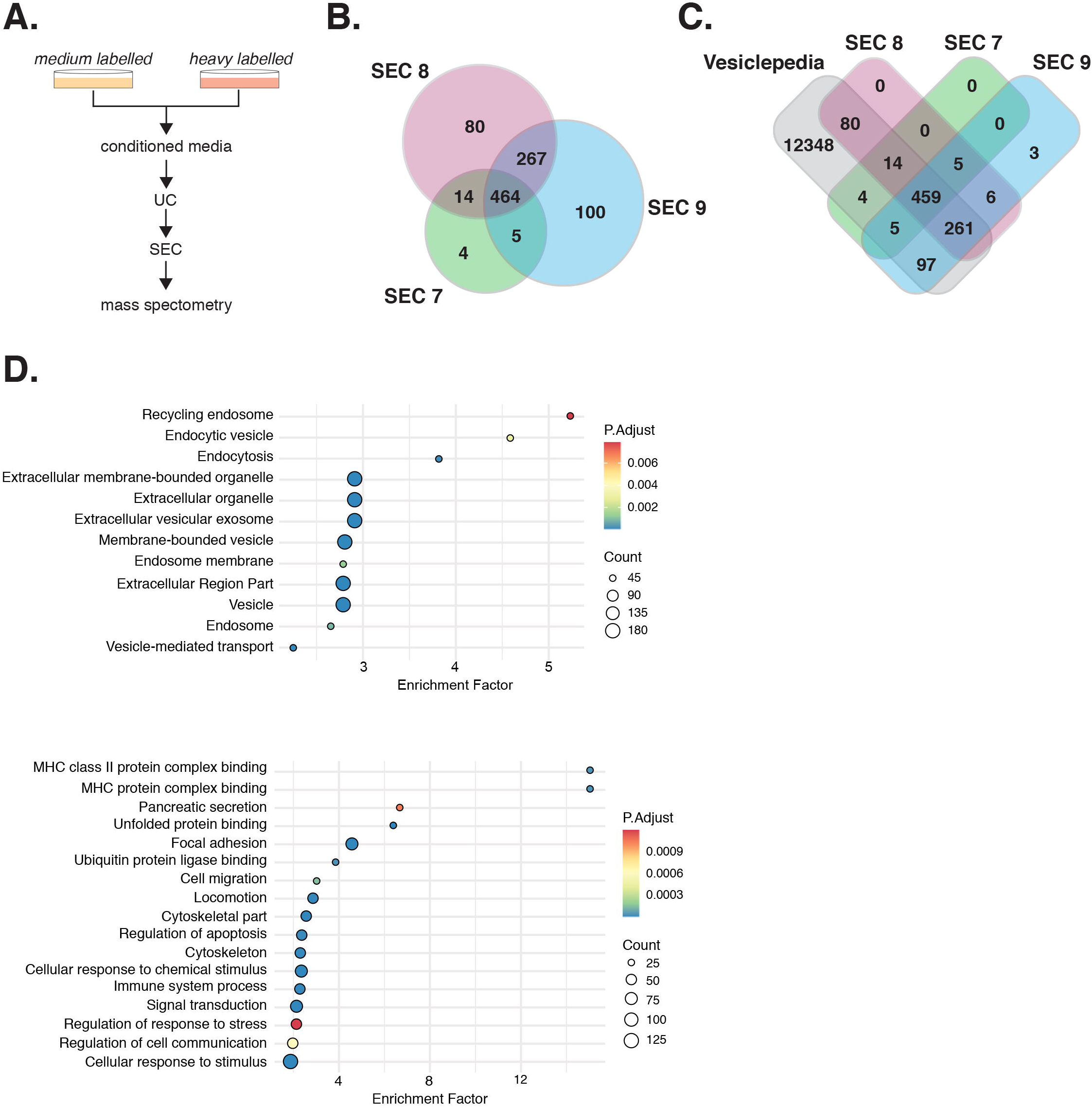
Proteomic analyses of _S_EVs secreted by cells with extra centrosomes support their endocytic origin. (A) Experimental flowchart. (B) Venn diagram comparing the _S_EVs proteomes of SEC fractions 7, 8 and 9. (C) Venn diagram comparing the depicting the _S_EVs proteome of SEC fractions 7, 8 and 9 with the Vesiclepedia database. (D) Dotplot representation of the enrichment analyses performed for the common proteins in all SEC fractions. Only proteins that were identified in both forward and reverse labelling experiments were considered for this analysis. See also Figure S2 and Tables S1-S3.

To investigate if centrosome amplification impacts on _S_EV protein composition, we analyzed changes in the ratio of proteins present in heavy and medium labelled _S_EV. Protein abundance was initially median normalized to ensure that heavy and medium intensities in each sample were equivalent. Interestingly, for proteins that a SILAC ratio could be calculated for, the ratio values did not significantly change in any SEC fraction (Figure S2C and Table S4), suggesting protein composition is largely unchanged in _S_EV secreted from cells with and without extra centrosomes. Overall, our SILAC data demonstrate that while extra centrosomes do not induce a major change in the _S_EVs protein composition, the content of these _S_EVs is consistent with an endocytic origin, indicating that this fraction is likely enriched for exosomes.

### Impaired lysosomal function in cells with extra centrosomes promotes _S_EVs secretion

MVBs are generally destined for degradation, by fusion with the lysosomal compartment, or are trafficked to the cell periphery where they fuse with the plasma membrane, resulting in exosome secretion [27, 33]. Lysosome dysfunction can shift the fate of MVBs targeted for degradation to fusion with plasma membrane, leading to increased _S_EVs secretion (Figure 3A) [34–36]. We demonstrated previously that centrosome amplification increases ROS [10], which can disrupt lysosomal function [37, 38]. Therefore, we hypothesized that defective lysosomal degradation of MVBs could lead to increased _S_EVs secretion in cells with amplified centrosomes (Figure 3A). To test this, we first assessed whether induction of centrosome amplification led to increased ROS production in PDAC cell lines. Indeed, induction of extra centrosomes increased ROS in both PaTu-S.iPLK4 and HPAF-II.iPLK4 cell lines, as measured by the ratio of reduced versus oxidized glutathione, where a decrease indicates higher ROS levels. Increased ROS can be abolished by treating cells with the ROS scavenger N-acetyl cysteine (NAC), while hydrogen peroxide (H_2_O_2_) is sufficient to increase ROS levels in these cells (Figures 3B and S3A). Using Magic Red fluorescence intensity to assess the function of the lysosomal cathepsin B protease [39], we found that cells with extra centrosomes have compromised lysosomal function. Treating cells with NAC prevented this defect, indicating that it was ROS dependent (Figures 3C, 3D and S3B). Furthermore, levels of LAMP1, a lysosomal marker, did not change in cells with extra centrosomes or in response to increased ROS (Figures S3C-3E), suggesting that ROS specifically impair lysosome function, consistent with their role in disrupting the integrity of lysosomal membranes [37]. Next, we analyzed _S_EVs secretion in response to ROS. These analyses revealed that whilst increased ROS were sufficient to increase _S_EVs secretion in PDAC cells, preventing higher ROS production in cells with amplified centrosomes, using NAC, abolished enhanced _S_EVs secretion (Figure 3E). These results suggest that compromised lysosome function in cells with amplified centrosomes leads to _S_EVs secretion. In agreement, inhibition of lysosome function with the vacuolar proton pump inhibitor Bafilomycin A1, which impairs lysosome acidification [40], was sufficient to increase _S_EVs secretion (Figures S3F-S3H) [41].

**Figure 3.**
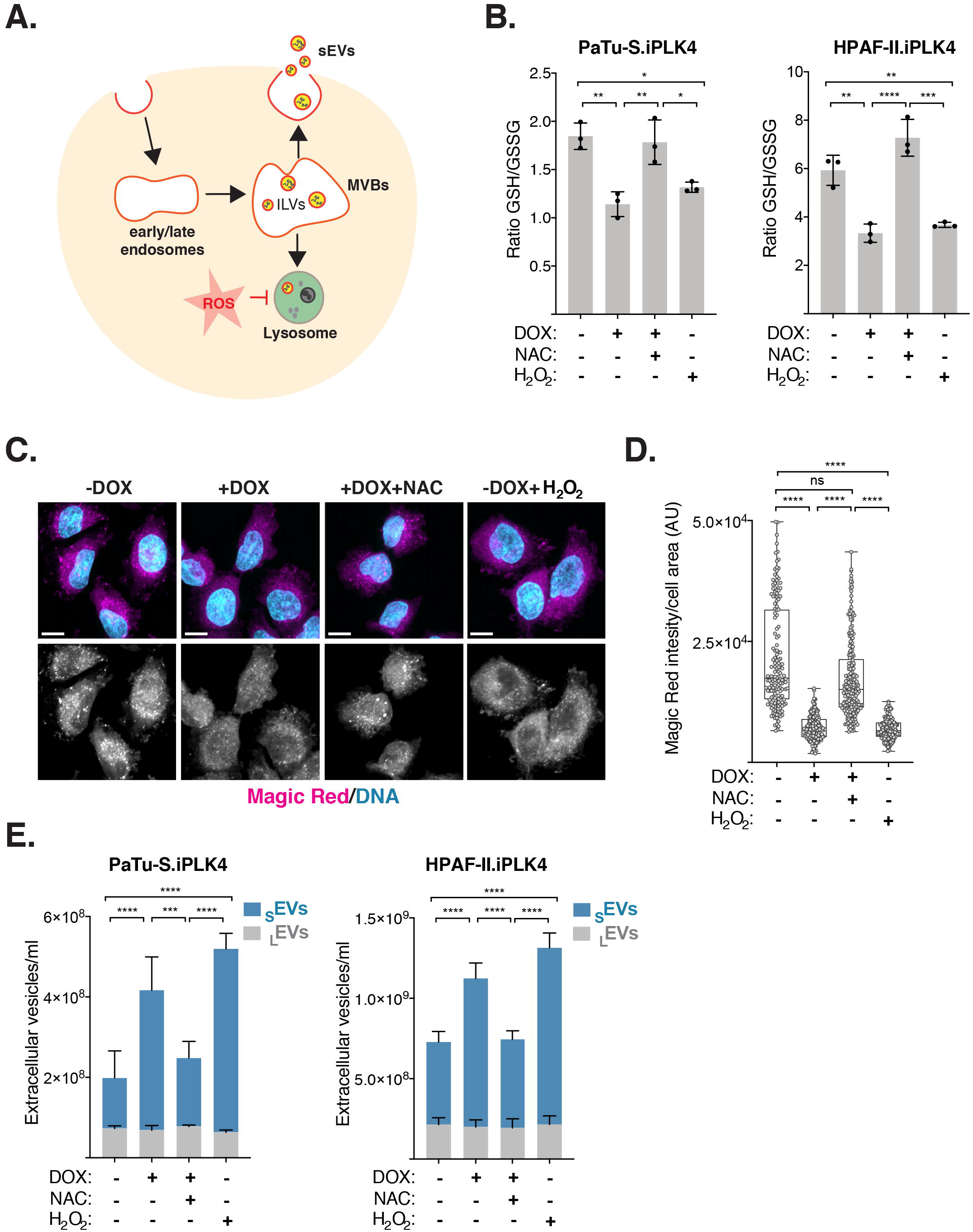
ROS promote lysosome dysfunction and _S_EVs secretion in cells with extra centrosomes. (A) Schematic representation of intraluminal vesicle formation (ILVs) and multivesicular bodies (MVBs) fate and how ROS could affect this process. (B) Levels of intracellular ROS quantified by the ratio of GSH/GSSG in Patu-S.iPLK4 and HPAF-II.iPLK4 cell lines. Decrease in the GSH/GSSG ratio indicates higher ROS levels. 5 mM of NAC and 100 μM H_2_O_2_ was used. (C) Representative confocal images of cells stained with Magic red (magenta), as a proxy for lysosome function, and for DNA (cyan). Scale bar, 10 μm. (D) Quantification of intracellular Magic red fluorescence intensity normalized for cell area in Patu-S.iPLK4 cells. AU, arbitrary units. 5 mM of NAC and 100 μM H_2_O_2_ was used. n_(−DOX)_=158, n_(+DOX)_=189, n_(+DOX+NAC)_=221 and n_(−DOX+H2O2)_=175. (E) Quantification of secreted _S_EVs and _L_EVs in Patu-S.iPLK4 and HPAF-II.iPLK4 cell lines. For all graphics error bars represent mean +/− SD from three independent experiments. \**p* < *0.05,* \*\**p* < *0.01*, \*\*\**p* < *0.001*, \*\*\*\**p* < *0.0001,* n.s. = not significant (*p* > *0.05*). The following statistic were applied: for graphs in B one-way ANOVA with Tukey’s post hoc test, for D one-way ANOVA with a Kruskal-Wallis post hoc test and for E two-way ANOVA with Tukey’s post hoc test. See also Figure S3.

Next, we investigated if ROS could prevent fusion between MVBs and lysosomes, thereby promoting MVB fusion with the plasma membrane and release of _S_EVs (Figure 3A). Using an antibody against phospholipid lysobisphosphatidic acid (LBPA), a lipid enriched at the membranes of late endosomes and MVBs [42], and lysotracker as a pH-based dye for functional lysosomes [43], we quantified the co-localization of MVBs and lysosomes in the different conditions. Centrosome amplification decreased the number of lysotracker-positive intracellular vesicles in a ROS-dependent manner but not LBPA-positive intracellular vesicles, further supporting defective lysosomal function as consequence of centrosome amplification (Figures 4A-4C). Strikingly, the percentage of co-localization between MVBs and lysosomes was significantly decreased in cells with extra centrosomes. NAC treatment restored lysosome function and MVB-lysosome co-localization in cells with extra centrosomes, while H_2_O_2_ was sufficient to decrease MVBs-lysosome co-localization (Figures 4A, 4B and 4D). Moreover, impairing lysosome function with Bafilomycin A1 dramatically reduced MVB-lysosome co-localization (Figures S4A-S4D). Taken together, our data suggest that decreased MVB-lysosome fusion as a consequence of increased ROS, and subsequent lysosome dysfunction, promotes _S_EVs secretion in cells with supernumerary centrosomes.

**Figure 4.**
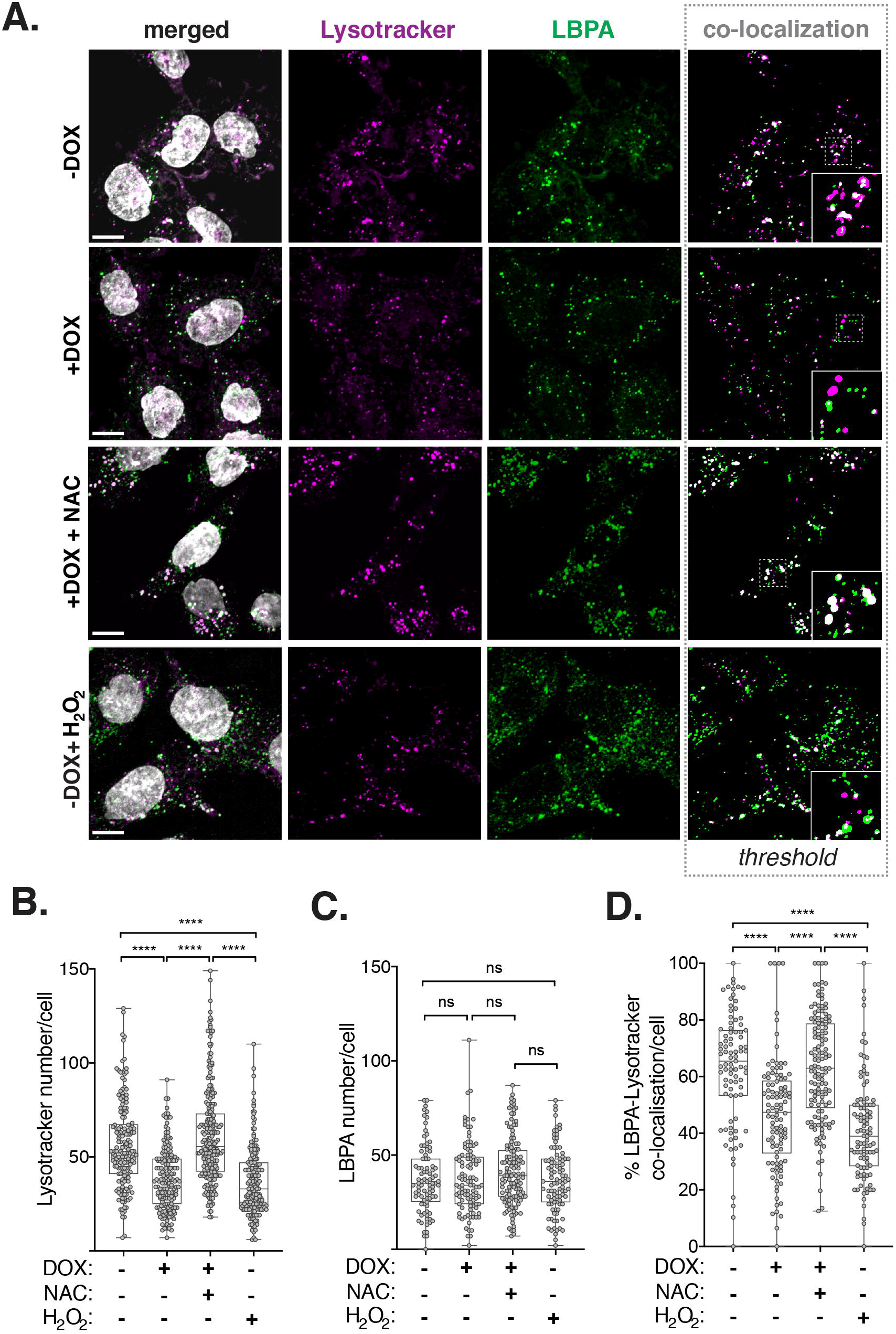
Centrosome amplification decreases lysosome-MVBs co-localization in a ROS-dependent manner. (A) Representative confocal images of cells stained for acidic lysosomes (Lysotracker, magenta), late endosomes/MVBs (anti-LBPA, green) and DNA (grey). Insets show higher magnification of lysotracker and LBPA-labelled vesicles. Scale bar, 10 μm. (B) Quantification of the number of lysotracker-labelled lysosomes per cell. 5 mM of NAC and 100 μM H_2_O_2_ was used. n_(−DOX)_=166, n_(+DOX)_=182, n_(+DOX+NAC)_=245 and n_(−DOX+H2O2)_=187. (C) Quantification of LBPA-labelled late endosomes/MVBs per cell. 5 mM of NAC and 100 μM H_2_O_2_ was used. n_(−DOX)_=88, n_(+DOX)_=102, n_(+DOX+NAC)_=129 and n_(−DOX+H2O2)_=x99. (D) Quantification of the percentage of lysotracker and LBPA-labelled intracellular vesicles co-localization. 5 mM of NAC and 100 μM H_2_O_2_ was used. n_(−DOX)_=86, n_(+DOX)_=102, n_(+DOX+NAC)_=129 and n_(−DOX+H2O2)_=98. For all graphics error bars represent mean +/− SD from three independent experiments. \*\*\*\**p* < *0.0001,* n.s. = not significant (*p* > *0.05*). For all graphs a one-way ANOVA with a Kruskal-Wallis post hoc test was applied. See also Figure S4.

### _S_EVs secreted by cells with extra centrosomes activate pancreatic stellate cells to facilitate cancer cell invasion

Cancer-associated _S_EVs often carry altered cargoes, rendering them functionally distinct from _S_EVs secreted by non-transformed cells [15, 16]. The exact causes of these changes, however, remain elusive. In PDAC, secreted _S_EVs may contribute to fibrosis through the activation of PSCs [44]. Thus, we investigated whether _S_EVs secreted by PDAC cells with extra centrosomes could promote the activation of PSCs. _S_EVs collected from PDAC cells −/+ extra centrosomes (donor cells) were added to PSCs (Figures 5A and 5B). Equal numbers of _S_EVs were added per condition to ensure that any differences observed were not due to the number of secreted _S_EVs. Activation of PSCs cells was assessed by immunofluorescence of alpha smooth muscle actin (αSMA). Increased expression and association of αSMA with stress fibers is a common feature of PSC activation towards a myofibroblast-like phenotype [45] (Figure 5C). Interestingly, treatment of PSCs with _S_EVs secreted by PDAC cells with extra centrosomes led to activation of ^~^25-30% of the cell population (Figure 5D). It is important to note that by normalizing _S_EVs numbers, we are likely underestimating the differences between _S_EVs secreted by cells −/+ centrosome amplification. As a positive control, PSCs were treated with TGF-β, a well-established activator of PSCs, known to lead to a strong activation phenotype (Figures S5A and S5B) [46]. To validate these results, we further purified the _S_EVs by SEC (Figures S2B and S5C) and tested the activation potential of the different isolated fractions. Not only were the _S_EVs harboring the potential to activate PSCs retained after SEC fractionation, but these _S_EVs associated mainly with one fraction (SEC8 for PaTu-S.iPLK4 and SEC9 for HPAF-II.iPLK4) (Figure 5E), further supporting the idea that secreted _S_EVs are indeed heterogeneous.

**Figure 5.**
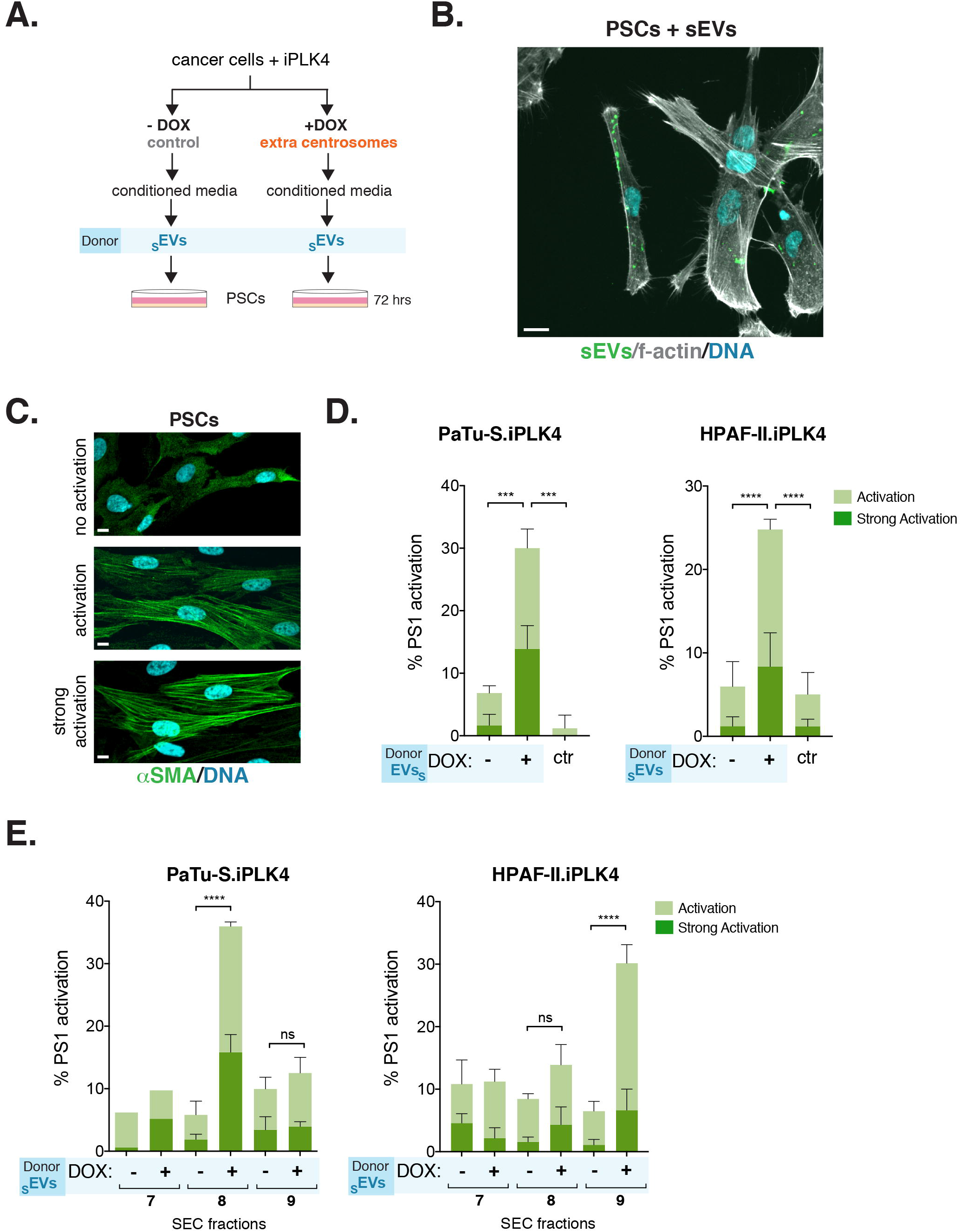
_S_EVs secreted by PDAC cells with amplified centrosomes activate pancreatic stellate cells. (A) Experimental flowchart. (B) Representative confocal image of PSCs incubated with _S_EVs. Cells were stained for f-actin (phalloidin, grey) and DNA (cyan). Isolated _S_EVs were labelled with Bodipy (green). Scale bar = 10 μm. (C) Representative confocal images of PSCs stained for α-SMA (green) and DNA (cyan). Scale bar, 10 μm. (D) Quantification of the percentage of PSCs activation upon treatment with _S_EVs collected by UC from Patu-S.iPLK4 (left) and HPAF-II.iPLK4 (right), with (+DOX) and without (−DOX) extra centrosomes. Patu-S.iPLK4 isolated SEV: PSCs n_(−DOX SEVs)_=398, n_(+DOX SEVs)_=373, and n_(ctr)_=475. HPAF-II.iPLK4 isolated _S_EV: PSCs n_(−DOX SEVs)_=914, n_(+DOX SEVs)_=1057, and n_(ctr)_=718. (E) Quantification of the percentage of PSCs activation upon treatment with SEVs collected by SEC from Patu-S.iPLK4 (left) and HPAF-II.iPLK4 (right), with (+DOX) and without (−DOX) extra centrosomes. Patu-S.iPLK4 isolated _S_EV: PSCs n_(−DOX SEVs SEC7)_=161, n_(+DOX SEVs SEC7)_=154, PSCs n_(−DOX SEVs SEC8)_=490, n_(+DOX SEVs SEC8)_=387, PSCs n_(−DOX SEVs SEC9)_=463, n_(+DOX SEVs SEC7)_=454. HPAF-II.iPLK4 isolated SEV: PSCs n_(−DOX SEVs SEC7)_=499, n_(+DOX SEVs SEC7)_=410, PSCs n_(−DOX SEVs SEC8)_=541, n_(+DOX SEVs SEC8)_=713, PSCs n_(−DOX SEVs SEC9)_=1035, n_(+DOX SEVs SEC7)_=914. For all graphics error bars represent mean +/− SD from three independent experiments. \*\*\**p* < 0.001, \*\*\*\**p* < *0.0001,* n.s. = not significant (*p* > *0.05*). For all graphs were analyzed using by two-way ANOVA with Tukey’s post hoc test. See also Figure S5.

Fibroblast activation is a common feature of cancer and can promote cancer cell invasion through various mechanisms including ECM remodeling and proteolysis [47]. To determine the functional relevance of PSC activation by _S_EVs secreted by PDAC cells with amplified centrosomes, we investigated their impact on PDAC cell invasion. To do so, we used 3-D heterotypic cultures of HPAF-II cells that form spheroids in 3-D with PSCs (Figure 6A) [48]. In contrast to non-treated PSCs, or PSCs pre-treated with _S_EVs from cells with normal centrosome numbers, PSCs pre-treated with _S_EVs harvested from cancer cells with extra centrosomes significantly induced invasion (Figures 6B and 6C). TGF-β pre-treated PSCs, used as positive control, showed higher invasion potential, consistent with the stronger levels of PSC activation observed (Figures 6B, 6C and S5B). Confocal imaging of 3-D spheroids composed of cancer cells expressing H_2_B-RFP and PSCs expressing H_2_B-GFP revealed that activated PSCs lead the invasive front (Figure 6D). Our findings demonstrate that _S_EVs secreted by PDAC cells with extra centrosomes are functionally different and can induce PSCs activation to promote cancer invasion.

**Figure 6.**
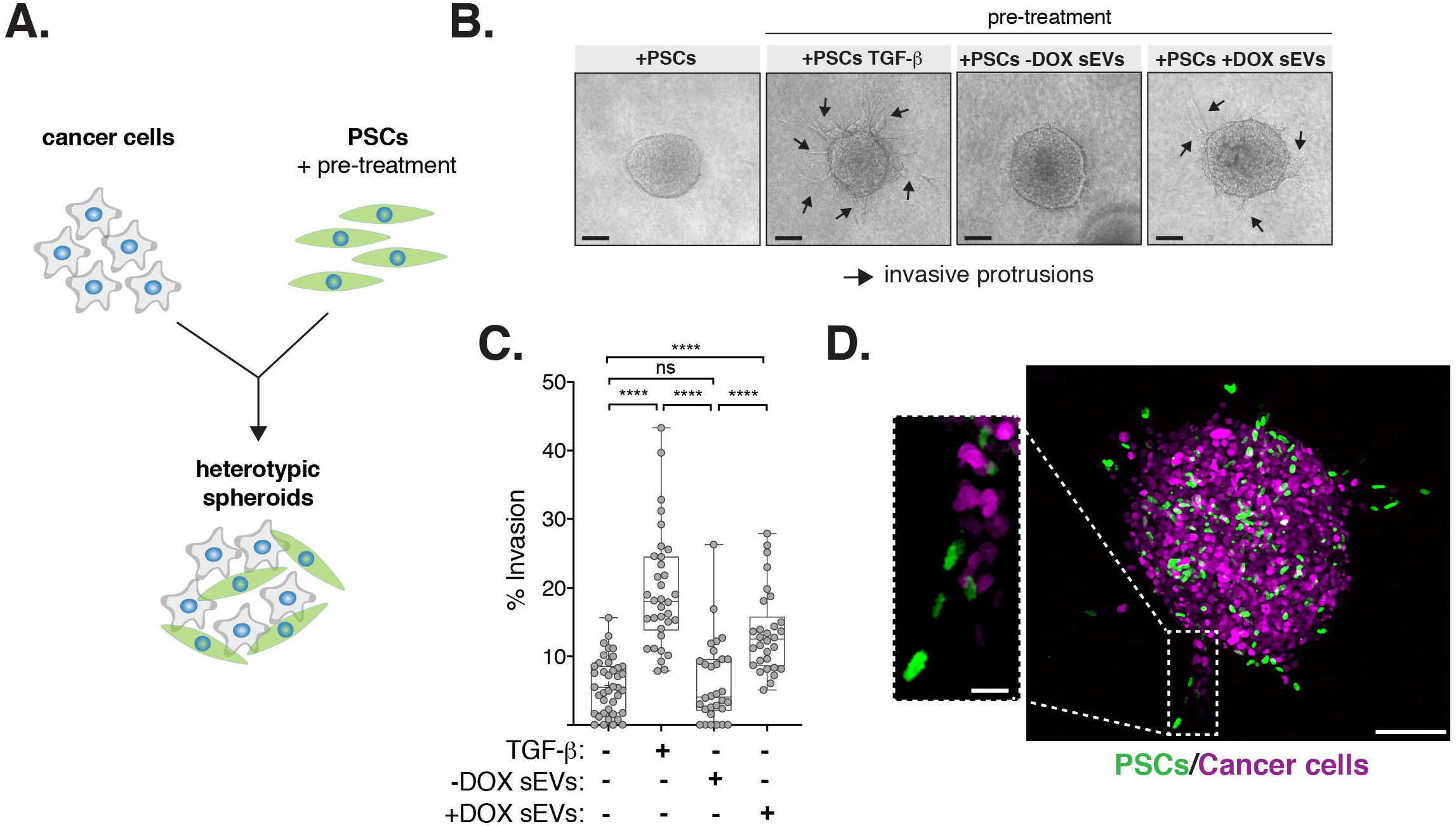
_S_EVs secreted by cells with extra centrosomes can promote PDAC invasion. (A) Experimental flowchart. (B) Representative brightfield images of heterotypic spheroids. Black arrows: invasive protrusions. Scale bar, 100 μm. (C) Quantification of the percentage of invasion in 3D spheroids. 5 ng/ml TGF-β was used as positive control. Spheroids n_(+PSCs)_=40, n_(+PSCs TGF-β)_=34, n_(+PSCs −DOX _S_EVs)_=31 and n_(+PSCs +DOX SEVs)_=31. (D) Confocal images of spheroids composed of cancer cells (expressing H_2_B-RFP; magenta) and PSCs (expressing H_2_B-GFP; green). Scale bar, 100 μm. Insect depicts higher magnification of invasive protrusion. Scale bar, 20 μm. For all graphics error bars represent mean +/− SD from three independent experiments. \*\*\*\**p* < *0.0001,* n.s. = not significant (*p* > *0.05*). Graph was analyzed using one-way ANOVA with a Kruskal-Wallis post hoc test.

## Discussion

In this study, we demonstrate that centrosome amplification induces secretion of _S_EVs that activate PSCs promoting the invasion of cancer spheroids. Activated PSCs are major players in the development of the pancreatic cancer stroma and associated fibrosis [21–23], suggesting a role for centrosome amplification in shaping the pancreatic cancer TME. Our data support a model whereby elevated ROS levels induced by extra centrosomes lead to loss of lysosomal function, favoring MVBs fusion with the plasma membrane and _S_EVs secretion (Figure 7).

**Figure 7.**
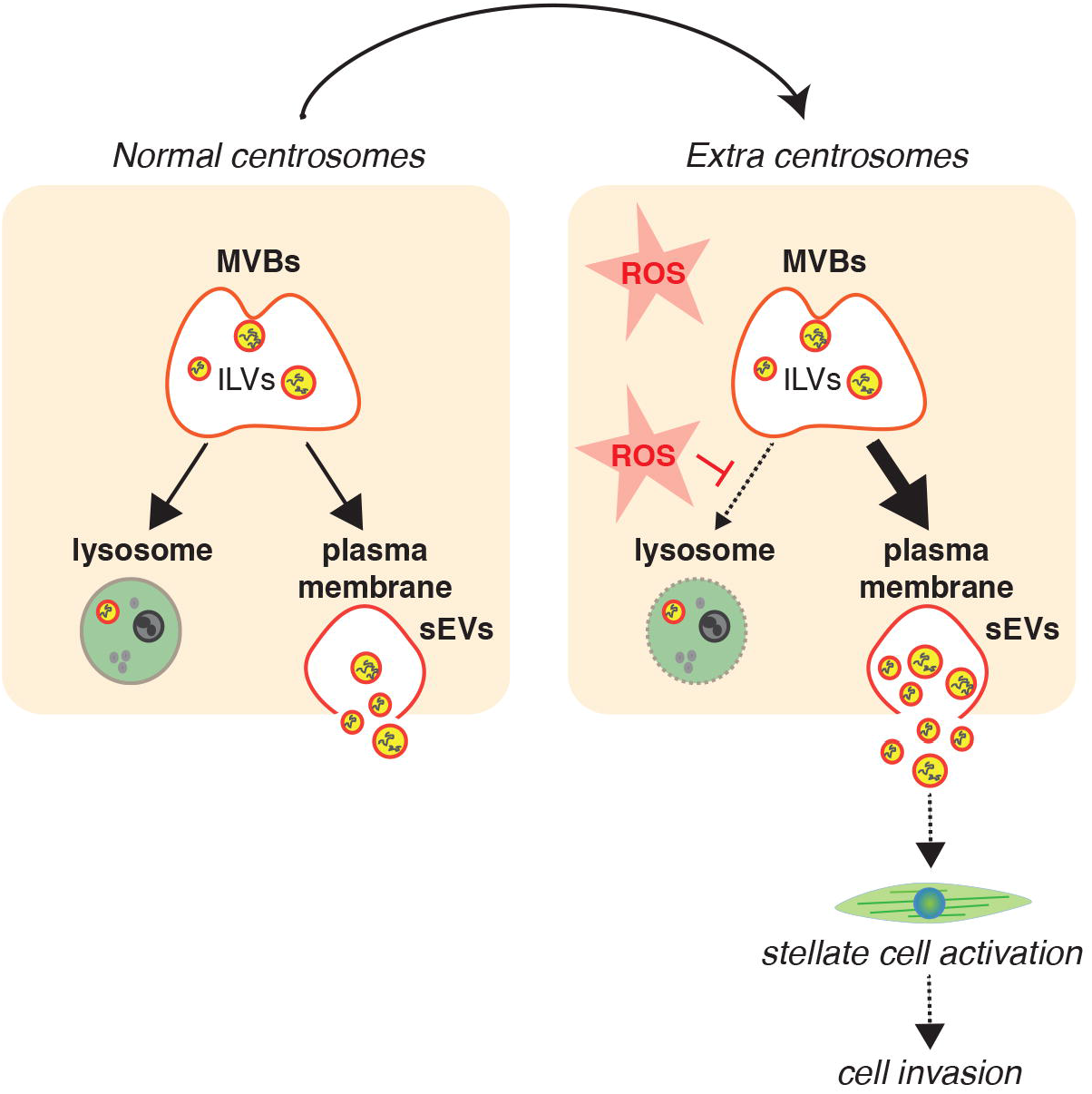
Schematic representation of working model. Increased ROS levels in cells with extra centrosomes compromises lysosomal function. We propose that this changes MVBs fate towards fusing with the plasma membrane and secretion of _S_EVs. _S_EVs secreted by cancer cells with extra centrosomes are functionally distinct and can induce PSCs activation to promote cell invasion.

Lysosomes are signaling centers that integrate many cellular responses to changes in nutrients, growth factors and stresses [49]. Fusion of lysosomes with autophagosomes is critical during autophagy, a self-degradative process important for the removal of protein aggregates, damaged organelles and intracellular pathogens [49]. Interestingly, centrosome amplification was recently shown to disrupt autophagy, rendering these cells sensitive to autophagy inhibitors [50]. Whether lysosome dysfunction is responsible for the autophagy defects observed in these cells is currently unknown. However, it is reasonable to assume that ROS-mediated lysosomal dysregulation could have a broader impact on the physiology of cells carrying centrosomal abnormalities.

_S_EVs secreted by cells with extra centrosomes exhibit many characteristics of exosomes: correct size range (30-150nm) and proteomic profiling revealed an enrichment for proteins associated with exosomes and exosome biogenesis. Sub-fractionation of secreted _S_EVs by SEC demonstrated not only the existence of different sub-types of _S_EVs, as previously reported [51, 52], but that functional differences between these different _S_EV populations also exist, as assessed by their ability to activate stellate cells. How changes in _S_EVs composition occur and how these induce stellate cell activation, however, remains elusive. One possibility is that changes in the _S_EV cargoes (proteins, RNA species) could be involved in stellate cell activation. Indeed, whilst SILAC ratio values for most detected proteins remained unchanged, we identified a number of proteins that were only identified in one label, for which a SILAC ratio could not be calculated. Therefore, it is possible that some of these proteins could play a role in PSC activation, but further studies will be required to assess if this is the case.

Alternatively, the presence or absence of specific proteins could influence _S_EV uptake and indirectly contribute to PSC activation. Cargo transfer by EVs can be mediated by delivery of surface proteins to membrane receptors, fusion with the plasma membrane, micropinocytosis, phagocytosis and receptor-mediated endocytosis to deliver their content [53]. In addition, interaction between EVs and secreted proteins has been shown to modulate their uptake, highlighting the complex regulation of this process [54]. Tetraspanins, such as CD9, CD63 and CD81, have been shown to be involved in the interplay between adhesion molecules and integrins to promote _S_EV uptake [55]. The presence of specific tetraspanins could also influence the specificity of target cells. For example, _S_EVs lacking the expression of the tetraspanin CD63 were found to be preferentially endocytosed by neurons [56]. Interestingly, we found that CD81 was the only protein absent specifically in the _S_EVs harvested from PDAC cells with amplified centrosomes that activate PSCs. Similarly, loss of CD81 has previously been reported in _S_EVs that are secreted upon induction of lysosome dysfunction [35]. Whilst the reason for this CD81 loss in response to lysosomal dysregulation is currently unknown, the striking similarity suggests a common response to lysosomal defects that could potentially modulate _S_EV uptake.

In summary, we describe a mechanism by which a stress response downstream of extra centrosomes culminates with the secretion of functionally different _S_EVs by diverging the fate of MVBs. Several cellular stresses have been shown to induce EV secretion, such as oxidative stress, hypoxia and radiation-induced cell stress [57]. Thus, it is possible that in response to multiple stressors, MVBs that are normally targeted for lysosomal degradation play a role in the release _S_EVs carrying protective functions in order to maintain tissue homeostasis. Indeed, oxidative stress itself has been shown to induce changes in the mRNA content of exosomes secreted by mouse mast cells, which help to protect the surrounding cells by conferring resistance to subsequent oxidative insult [58]. Understanding how stress communication protects cancer cells could allow us to exploit these mechanisms to prevent cancer cell adaptation.

## Acknowledgements

We are grateful to Judith Simon and all the members of the Godinho lab for comments or discussion of the manuscript; Andrea Lafuente for providing the Dotplots used in Figure 2D; Hefin Rhys for helping with ImageStream and Nanosight; Pedro Monteiro for advice on image analyzes. This work was supported by a Cancer Research UK Centre Grant to Barts Cancer Institute (C355/A25137). E.P.C. and R.G. were funded by Cancer Research UK (C10847/A27781). G.R. is supported by the French Government (ANR) through the 20 Investments for the Future LABEX SIGNALIFE (ANR-11-LABX-0028-01) and by the Fondation pour la Recherche Medicale (DEQ2011104211324). F.M. is supported by a Medical Research Council (MRC) Career Development Award (MR/P009417/1) and a Barts Charity grant (MGU0346). S.D.A. was supported by an MRC PhD studentship and J.C. supported by a Barry Reed PhD studentship. S.A.G. is a fellow of the Lister Institute and is supported by the MRC (MR/T000538/1).

## Author Contributions

Conceptualization: S.D.A., S.A.G.; Methodology: S.D.A., T.A., G.D., E.P.C., R.G., G.R., F.M., S.A.G.; Validation: S.D.A., J.C.; Formal Analysis: S.D.A., J.C., G.D., F.M., S.A.G.; Investigation: S.D.A., J.C., T.A., G.D., E.P.C., M.D.; Resources: H.K., R.G., G.R., F.M., S.A.G.; Data Curation: S.D.A., J.C., F.M.; Writing – Original Draft: S.A.G.; Writing – Review & Editing: S.A.D., G.D., E.P.C., H.K., R.G., G.R, F.M., S.A.G.; Visualization: S.A.D., J.C., G.D., E.P.C., S.A.G.; Supervision: H.K., G.R., F.M., S.A.G.; Project Administration: S.A.G.; Funding Acquisition: S.A.G.

## Methods

### Cell culture

Adherent cell lines were cultured at 37°C and 5% humidified CO_2_. The pancreatic cancer cell lines PaTu-8988t (PaTu-T; gift from Y. Wang, BCI-QMUL) PaTu-8988s (PaTu-S), Capan-1, PANC-1, CFPAC-1, HPAF-II, MIA-PaCa-2 and DEC-hTERT (derived from normal pancreas) (gifts from H. Kocher, BCI-QMUL) were grown in DMEM supplemented with 10% FBS and 1% penicillin and streptomycin. HPDE cells (derived from normal pancreas) (gift from H. Kocher, BCI-QMUL) were grown in keratinocyte-SFM (1X) serum free media +30μg/ml (BPE)+ 0.2ng/ml rEGF. The pancreatic stellate cell lines PS1 (gift from H. Kocher, BCI-QMUL) [59] were grown in DMEM/F12 supplemented with 10% FBS and 1% penicillin and streptomycin. 5 ng/ml of recombinant TGF-β (Peprotech) was used to treat PS1 cells for 72 hours. Tetracycline-free FBS was used to grow cells expressing the PLK4 Tet-inducible construct. STR profiling was performed for cell line authentication on the following lines: PaTu-S, PaTu-T, Capan-1, MIA-PaCa-2, Panc-1 and PS1.

### Chemicals

Chemicals and treatments were performed as follows: 2μg/ml Doxycycline hyclate (DOX; Sigma) for 48 hours, 100 μM H_2_O_2_ (Sigma) for 48 hours, 5 mM N-acetyl cysteine (NAC; Sigma) for 48 hours and 20 nM Bafilomycin A1 (Sigma) for 24 hours.

### Lentiviral production and Infection

To generate lentivirus, HEK-293 cells were plated in antibiotic free medium. Transfection of the appropriate lentiviral plasmid in combination with Gag-Pol (psPAX2, Addgene, 12260) and VSV-G (VSV-G: pMD2.G, Addgene, 12259) was performed using lipofectamine 2000^®^ (Thermo Fisher Scientific), as per the manufacturer’s specifications. The resultant lentivirus was harvested 24 hours and 48 hours post infection, passed through a 0.4 μM syringe filter and stored in cryovials at −80°C. For infection, the appropriate lentivirus was then mixed with 8 μg/ml polybrene before being added to the cells in a dropwise fashion. Infection was repeated the following day and antibiotic selection started 24 hours after final infection.

Cells expressing the inducible PLK4 construct were generated as previously described [26]. Briefly, cells were initially infected with pLenti-CMV-TetR-Blast lentiviral vector (Addgene, 17492) and selected using Blasticidin (10 μg/ml). Post-selection, cells were then infected with a lentiviral vector containing PLK4 cDNA which had been previously cloned into the pLenti-CMV/TO-Neo-Dest vector and selected using Geneticin (200 μg/ml) [26, 60]. Cells expressing the PLK4 transgene were then induced for 48 hours using 2 μg/ml of Doxycycline.

To generate H2B-RFP iPLK4 cells, lentivirus was produced by transfecting HEK-293 cells with LV-RFP (Addgene 26001), pMD2.G (Addgene, 12259) and μg pCMVDR8.2 (Addgene, 12263) using FuGENE (Promega, E2311), as per manufacturer’s instructions. 24 hours later the medium was replaced and 48 hours post transfection the viral supernatant was collected, passed through a 0.4 μM syringe filter and stored in cryovials at −80°C. Cells were transduced with the lentivirus as described above.

### siRNA

siRNA transfection was performed in antibiotic free growth medium using Lipofectamine^®^ RNAiMAX as per the manufacturer’s specifications. For SAS-6 knock down experiments siNegative control (siNegative, Qiagen, 1027310) and siSAS-6 (siSAS6 on-TARGET smart pool, Dharmacon, M-004158-02) were used. Per 6 well, 20 nM of siRNA was used for PaTu-S.iPLK4 cells and 50 nM for HPAF-II.iPLK4 cells as PaTu-S.iPLK4 cells were more sensitive to SAS-6 depletion and to prevent loss of centrioles below control conditions. 24 hours post transfection, the cells were trypsinized and seeded onto coverslips for analysis by immunofluorescence or into 15 cm dishes for exosome harvest experiments 72 hours post transfection.

### Immunofluorescence 2D

Cells plated on glass coverslips were treated for up to 48 hours with the appropriate drug treatments, before being washed twice in PBS and fixed in 4% Formaldehyde for 20 minutes at room temperature. For centrin2 staining, cells were fixed in ice-cold methanol for 10 minutes at −20 ° C. Following fixation, cells were permeabilized in 0.2% Triton X-100 in PBS for 5 minutes then blocked for 30 minutes in blocking buffer (PBS, 5% BSA, 0.1% Triton X-100). Cells were then incubated with primary antibody diluted in blocking solution for 1 hour. Cells were then washed with PBS and incubated with species-specific Alexa-conjugated secondary antibodies diluted in blocking buffer for 1 hour. Alexa Fluor 568 Phalloidin (1:250) was incubated in blocking solution for 1 hour. Cells were washed in PBS and DNA was stained with Hoechst 33342 diluted in PBS (1:5000) for 5 minutes. Finally, coverslips were mounted using ProLong™ Gold Antifade Mountant. Antibodies used included: Anti-centrin 2 N-17-R (Santa Cruz; 1:100), Anti α-tubulin DM1 α (Sigma-Aldrich; 1:1000), Anti LBPA 6C4 (Merck Millipore; 1:100), Anti LC3B (D11) XP ^®^ (Cell Signalling; 1:200), Anti α-SMA (Sigma-Aldrich; 1:300), Anti-Rabbit Alexa Flour 488 (Life Technologies; 1:1000), Anti-Rabbit Alexa Fluor 568 (Life Technologies 1:1000), Anti-Mouse Alexa Fluor 488 (Life Technologies 1:1000). Centrosome amplification was defined as the percentage of metaphase cells containing extra centrosomes (>4 centrioles per cell).

Images were acquired using an inverted Nikon microscope coupled with a spinning disk confocal head (Andor). Unless otherwise stated, imaging of cancer cells was performed using a 100x objective and imaging of stellate cells with a 40x objective. Images/projection images (from z-stacks) were subsequently generated and analyzed with Image J (National institute of Health, Bethesda, MD, USA) [61]. Where Z-stack images were required to analyze fluorescence intensity, Z-stack parameters were determined using the following equation: Zmin = 1.4λn/(NAobj)2. λ = the emission wavelength, n= refractive index of the immersion media, NAobj = the numerical aperture of the objective. This equation calculates the ideal z stack step size to minimize overlap between each step of the stack. Sum intensity projection images were subsequently generated using Image J and fluorescence intensity was quantified using Image J. All conditions were quantified blindly.

### Extracellular Vesicle (EV) Isolation

Cells were grown for 48 hours in medium supplemented with EV depleted FBS. Vesicle depletion in FBS was performed via ultracentrifugation at 100,000 x g at 4°C for 18 hours. Where induction of centrosome amplification was necessary, cells were treated with DOX for 48 hours, before cells were washed in PBS and subsequently cultured in EV depleted media. Where drug treatments were required, cells were treated for the duration of the EV harvest (48 hours post addition of EV depleted media). After 48 hours, conditioned medium was collected, and a final cell count was performed to ensure the final cell count remained the same between cell types and conditions.

#### Serial ultracentrifugation (UC)

Extracellular vesicles were isolated from the conditioned media via serial ultracentrifugation steps at 4°C, similarly to [14]. Briefly, the cell culture medium was subjected to a low speed centrifugation of 500 x g for 10 minutes. The supernatant was then centrifuged at 12,000 x g for 20 minutes to pellet the large EVs (_L_EVs), after removal of the supernatant the _L_EVs were re-suspended in 500μl of PBS. The supernatant was then subjected to a high-speed ultracentrifugation at 100,000 x g for 70 minutes to pellet the smaller EVs (_S_EVs). The pellet was then washed in PBS and a second high-speed ultracentrifugation was performed at 100,000 x g for 70 minutes (Figure S1A). The isolated _S_EV pellet was then re-suspended in 500 μl of PBS.

#### Size exclusion chromatography (SEC)

To further purify EVs isolated by serial ultracentrifugation, size exclusion chromatography (SEC) was performed using the qEV original izon science SEC columns (as per the manufacturer’s instructions). Briefly, the SEC columns were equilibrated to room temperature and flushed with 5ml of buffer (PBS filtered twice through 0.22 μM filters) prior to use. 500 μl of concentrated exosomes (isolated by serial ultracentrifugation) was added to the top of the column and the eluted fractions were collected immediately in 500 μl volumes. The column was kept topped up with buffer throughout the experiment. Fractions 7-12 containing the eluted EVs were collected.

### Extracellular Vesicle Quantification and Analysis

#### Amins ImageStream^®^ Mark II Imaging Flow Cytometer (ImageStream)

EV samples were analyzed by ImageStream as previously described [25]. Briefly, samples were prepared in 50 μl volumes, labelled with the fluorescent lipid dye BODIPY^®^ FL Maleimide [BODIPY^®^ FL N-(2-Aminoethyl) Maleimide] (Thermo Fisher Scientific; 1:200) and incubated at room temperature in the dark 10 minutes prior to analysis. Samples were then loaded onto the ImageStream and vesicles were acquired at a slow flow rate with 60x magnification, a 488 nm excitation laser (BODIPY detection) and 765 nm laser (side scatter). The “remove bead” function was turned off and the flow rate allowed to stabilize before acquisitions. For acquisition, the storage gate was set to collect all events and the stopping gate set to the vesicle population (low to mid BODIPY intensity and low side scatter). The stopping gate was set to ensure that at least 20,000 objects were analyzed per acquisition. Three separate acquisitions were collected per sample. Analysis was then performed using the IDEAS software. To quantify objects/ml, a graph was generated plotting channel 02 fluorescence intensity (BODIPY) against channel 12 scatter intensity (side scatter) and a vesicle gate was re-applied to select the population at the correct BODIPY and side scatter intensities to be EVs (see Figure S1C). Where necessary the gate was adjusted using the Image library to eliminate noise and artefacts from the vesicle population. The objects/ml statistic was then used to quantify the number of objects/ml in the gated region. The average objects/ml was calculated from three separate acquisitions from each sample.

#### Nanoparticle tracking anlaysis

Performed using a NanoSight NS300 with a high sensitivity camera and a syringe pump. As previously described, isolated EVs were resuspended (UC) or eluted (SEC) in Dulbecco’s PBS filtered twice through 0.22 μM filters. The NS300 chamber was flushed with 0.22 μM filtered deionized water and then again with 500 μl of PBS (Dulbecco’s PBS filtered twice through 0.22 μM filters) to remove any particle matter. Using a 1 ml syringe 400 μl of EV sample was then flushed through the chamber until vesicles were visible on the camera to allow the focus and gain settings to be optimized. The sample was then injected into the flow cell at speed 50 and 3 recordings of 60 seconds each were acquired. Between samples filtered PBS was used again to flush the chamber ensuring no residual particles remained. The data was then analyzed using the NTA 3.2 analysis software and averages of the three technical replicates were plotted per experiment.

### Immunolabeling electron microscopy (IEM)

A drop (5μl) of _S_EVs (isolated by UC) suspended in PBS was deposited on formvar-carbon-coated electron microscopy grids for 20 min at room temperature, fixed with 2% paraformaldehyde in 0.2 M phosphate buffer (pH 7.4), for 20 min at room temperature, and post fixed with 1% glutaraldehyde in PBS for 5 min at room temperature. Grids containing sEV were then washed and then blocked for 5 min at room temperature in blocking buffer (PBS, 1% BSA). _S_EVs were then immunolabelled with a mouse anti-human CD63 primary antibody (Abcam ab23792) diluted in blocking solution for 1 hour at room temperature, washed with PBS, 0,1 % BSA, incubated with a rabbit antibody against mouse Fc fragment (Dako Agilent Z0412) in PBS 0,1% BSA for 20 min at room temperature. The preparations were then immunogold labeled with protein-A gold-conjugates (10 nm; Cell Microscopy Center, Department of Cell Biology, Utrecht University). Grids were analyzed on a Tecnai Spirit G2 electron microscope (Thermo Fischer Scientific) and digital acquisitions were made with a 4k CCD camera (Quemesa, Soft Imaging System). Images were analysed with iTEM software (iTEM CE Olympus serie) and data with Prism-GraphPad Prims software (v8) [62].

### Western Blotting

Small extracellular vesicles harvested for protein extraction were isolated as previously described, however following the final ultracentrifugation, the pellet was lysed immediately in RIPA buffer supplemented with protease inhibitors on ice. To facilitate further lysis, samples were probe sonicated on ice. Protein concentration was determined using the Bio-Rad Protein Assay. 10μg of protein was loaded per well. Samples were resuspended in Laemmli buffer, resolved using the NuPAGE^®^ Bis-Tris Electrophoresis System with NuPAGE™ 10% Bis-Tris Protein Gels and transferred onto PDVF membranes. Antibodies used included: Anti TSG101 EPR7130(b) (Abcam; 1:1000), Anti CD63 (Abcam; 1:1000), Anti CD81 B-11 (Santa Cruz; 1:500), Anti ALIX 3A9 (Cell Signalling; 1:1000), Anti Flotillin-1 18 (Biosciences; 1:1000), HRP-anti rabbit secondary (GE Healthcare; 1:5000) and HRP-anti mouse secondary (GE Healthcare; 1:5000). Western blots were developed on X-ray film using a SRX-101A table top film processor.

### Stable isotype labelling by amino acids in cell culture (SILAC)

SILAC based proteomic analysis of exosomes was performed as previously [63]. All SILAC amino acids (heavy and medium) were purchased from Cambridge Isotopes. SILAC media and dialyzed serum were purchased from Thermo Fisher Scientific. PaTu-S.iPLK4 cells with and without the induction of centrosome amplification were grown for 6 passages in Dulbecco’s modified Eagle’s medium for SILAC supplemented with 10% Gibco ™ Dialyzed Fetal Bovine Serum (ultracentrifuged for 18 hours at 100,000 x g for EV depletion), 600 mg/L Proline and 100 mg/L of either heavy or medium Lysine and Arginine amino acids (Lys^8^ and Arg^10^ for heavy, and Lys^4^ and Arg^6^ for medium, respectively). Labelled cells were then plated at a density of 1×10^6^ cells in 40 T175 flasks per condition. 24 hours later flasks were washed in PBS and 15 ml of fresh EV depleted medium supplemented with the correct amino acids (heavy or medium) was added to the cells. 48 hours later, the conditioned medium was harvested and samples heavy and medium labelled were pooled together (Figure 2A). EVs were then isolated from the conditioned medium via ultracentrifugation and subsequent SEC as previously described. The experiment was then repeated with the labelling reversed.

### Mass spectrometry

Extracellular vesicles were lysed in 8 M Urea in 50 mM Ammonium bi-carbonate (ABC) (pH 8). Samples were then sonicated using a Diagenode Bioruptor sonicator at 4°C. Samples were sonicated at high power for 15 cycles of 30 seconds on and 30 seconds off. 10 mM DTT was added for 20 minutes at room temperature followed by 55 mM Iodoacetamide incubated for 30 minutes in the dark. Protein quantification was then performed as previously described. 15 μg of protein was then selected per sample and Urea was diluted to 2 M final concentration with 50 mM ABC. Samples were then subjected to in-solution trypsin digestion overnight at 25°C. The digested peptides were then acidified and desalted via stagetipping [64]. Peptides were then dried by vacuum centrifugation and resuspended in 10 μl of buffer A* (2% ACN, 0.1% trifluoroacetic acid and 0.5% acetic acid).

### LC-MS/MS analysis

Equivalent of ^~^1 μg of each digested SILAC mix was subjected to Liquid Chromatography coupled with tandem Mass Spectrometry (LC-MS/MS), using a Q-Exactive plus Orbitrap mass spectrometer coupled with a nanoflow ultimate 3000 RSL nano HPLC platform (Thermo Fisher Scientific). Briefly, samples were resolved at a flow rate of 250 nL/min on an Easy-Spray 50 cm × 75 μm RSLC C18 column with 2 μm particle size (Thermo Fisher Scientific), using a 123 minutes gradient of 3% to 35% of buffer-B (0.1% formic acid in ACN) against buffer-A (0.1% formic acid in water), and the separated peptides were infused into the mass spectrometer by electrospray. The spray voltage was set at 1.95 kV and the capillary temperature was set to 255 °C. The mass spectrometer was operated in data dependent positive mode, with 1 MS scan followed by 15 MS/MS scans (top 15 method). The scans were acquired in the mass analyzer at 375-1500 m/z range, with a resolution of 70,000 for the MS and 17,500 for the MS/MS scans. Fragmented peaks were dynamically excluded for 30 seconds.

### Proteomics data analysis

MaxQuant (version 1.6.3.3) software was used for database search and SILAC quantifications [65]. The search was performed against a FASTA file of the Homo Sapiens, extracted from Uniprot.org (2016). A precursor mass tolerance of 4.5 ppm, and a fragment mass tolerance of 20 ppm was applied. Methionine oxidation and N-terminal acetylation were included as variable modifications whilst carbamidomethylation was applied as a fixed modification. Two trypsin miss-cleavages were allowed, and the minimum peptide length was set to 7 amino acids. SILAC multiplicity was set to 3, with Lys4 and Arg6 selected as medium, and Lys8 and Arg10 as heavy labels. Minimum SILAC ratio count was set at 1. All raw files were searched together, with the match between runs option enabled. All downstream data analysis was performed by Perseus (version 1.5.5.3) [52], using the MaxQuant ProteinGroups.txt output file. Briefly, normalized SILAC H/M intensities were converted to Log 2 scale. Reverse (decoy) hits, potential contaminants, and proteins identified only by modified peptides were filtered out. Ratio values were then median subtracted. Category enrichment analysis was performed using the Fisher exact test function within Perseus. Scatter plots of the SILAC ratio values were also generated by Perseus. All mass spectrometry raw files and search results reported in this paper have been deposited at the ProteomeXchange Consortium via the PRIDE [66], with the PRIDE accession number of PXD020984.

### Measuring cellular reactive oxygen species (ROS)

Cellular ROS was measured through the detection glutathione in its reduced (GSH) and oxidized (GSSG) forms using the luminescence-based GSH/GSSG-Glo™ Assay (Promega, V6611). Briefly, the Promega GSH/GSSG-Glo™ Assay is a linked assay utilizing glutathione S-transferase and Luciferin-NT that generates a luminescent signal in response to levels of GSH present in the sample. The ratio of GSH to GSSG can then be calculated to give a read out of oxidative stress in the cells, where a decrease in the ratio indicates an increase in oxidative stress. All reactions and calculations were carried out as per the manufacturer’s instructions. The final ratio of GSH/GSSG was normalized to protein content to control for any changes in cell number. Protein was quantified using the Pierce™ BCA Protein Assay Kit (Thermo Fisher Scientific, 23227) as per the manufacturer’s instructions.

### Magic Red assay

The Magic Red™ Cathepsin B kit (Bio-Rad, ICT937) was used to analyze the protease activity of Cathepsin B in lysosomes as a proxy to lysosome function. In the presence of functional cathepsin B, the Magic Red substrate is cleaved allowing the Cresyl violet fluorophore to fluoresce red upon excitation at 550-590 nm. Briefly, cells to be analyzed were plated on coverslips and the Magic Red substrate (Magic Red stock was reconstituted in 50 μl DMSO and diluted 1:10 in deionized water) was added to the growth media (20μl was added per 300μl of growth media as per the manufacturer’s instructions) for the final hour of the experiment. Cells were then fixed in 4% Formaldehyde as previously described. Cresyl Violet fluorescence was detected using an inverted Nikon microscope coupled with a spinning disk confocal head (Andor). Z-stack images were acquired, and sum intensity image projections were generated using Image J. Fluorescence intensity was then quantified per cell with ImageJ [61]. All conditions were quantified blindly.

### Extracellular vesicle uptake by recipient cells

Fluorescently labelled EVs were generated using the previously described ultracentrifugation protocol with the following alteration: prior to the final PBS wash step, EVs were resuspended in 200 μl of PBS and fluorescently labelled with BODIPY (1:200). EVs were then incubated at room temperature for 5 minutes before being diluted in 31.5 ml of PBS. The final 100,000 x g ultracentrifugation step was then performed, and the subsequent EV pellet resuspended in 200 μl of PBS. The isolated EVs were then added to the recipient cells that had been plated on glass coverslips 24 hours prior. 3 hours post addition of EVs, coverslips were fixed in 4% formaldehyde and stained with Alexa Fluor 568 Phalloidin (1:250) and Hoechst (1:5000) as previously described. Representative z-stack images were taken using a spinning disk confocal microscope as previously described.

### Extracellular vesicle-mediated PSC activation assay

PaTu-S.iPLK4 cells untreated or induced to have amplified centrosomes (48 hours 2 μg/ml DOX treatment) were cultured for 48 hours in vesicle depleted media before the conditioned media was collected. EVs were then harvested from the conditioned media by ultracentrifugation alone, or in combination with SEC as described previously. EV number was then quantified by ImageStream as described above. 20 million EVs were then added to the culture medium of PS1 cells that had been plated on glass coverslips at a density of 1×10^4^ cells 24 hours prior. 48 hours later, a second dose of 20 million EVs was administered. 24 hours later cells were fixed and stained for α-SMA and DNA as described previously. Images were acquired using an inverted Nikon microscope coupled with a spinning disk confocal head (Andor) with a 40x objective. PS1 activation was quantified based on α-SMA organization, where the formation of α-SMA fibers was used as a measure of activation (maybe describe how the different categories were separated and we can reference the figure). Roughly 150 cells were quantified manually per condition. All conditions were quantified blindly.

### 3D co-culture spheroid invasion assay

Prior to spheroid generation, PS1 cells were either treated for 72 hours with _S_EVs (as described above), with 5ng/ml TGF-β or left untreated. 3D spheroid cancer cell/PS1 co-cultures were generated using a hanging drop spheroid model developed by Ed Carter and Richard Grose (BCI-QMUL), based on previous work [48]. Briefly, PS1 H2B-GFP (4.4× 10^4^ cells/ml) and HPAF-II.iPLK4-H2B-RFP cancer cells (2.2×10^4^ cells/ml) were combined in a 0.24% methylcellulose solution (Sigma-Aldrich, M0512). Droplets containing 1000 cells were then plated on the underside of a 15 cm culture dish and left to form spheroids overnight at 37 °C. Spheroids were then collected and centrifuged at 100x g for 3 minutes before being re-suspended in gel mix solution. Gel mix solution consisted of 1.6 mg/ml Collagen I (Corning Rat Tail High Concentration) and 17.5 % Matrigel ^®^ Matrix Basement Membrane LDEV-free (Corning, 354234), prepared in PS1 culture medium and buffered to physiological pH with NaOH. Approximately 6 spheroids suspended in gel mix were added to a pre-coated well of a low attachment plate and left to solidify at 37 °C before PS1 culture medium was added on top. Spheroids were incubated for 3 days and images were taken by light microscopy. Percentage invasion was analyzed using Image J and calculated as a measure of the total invasive area relative to the central sphere. For confocal analyses, spheres were fixed in 4% formaldehyde prior to mounting for imaging on an LSM 880 Zeiss confocal microscope. All conditions were quantified blindly.

### Statistical Analysis

Graphs and statistics were generated using Prism 8 (GraphPad Software) where results are presented as mean ± standard deviation (SD) unless otherwise stated. Statistical analysis was performed using one-way ANOVA with either a Tukey’s (parametric) or Kruskal-Wallis (non-parametric) post hoc test unless otherwise stated. Significance is equal to *p<0.05, **p<0.01, ***p<0.001 and ***P<0.0001.

