## Supplemental Figures for "Centrosome amplification mediates small extracellular vesicles secretion via lysosome disruption"

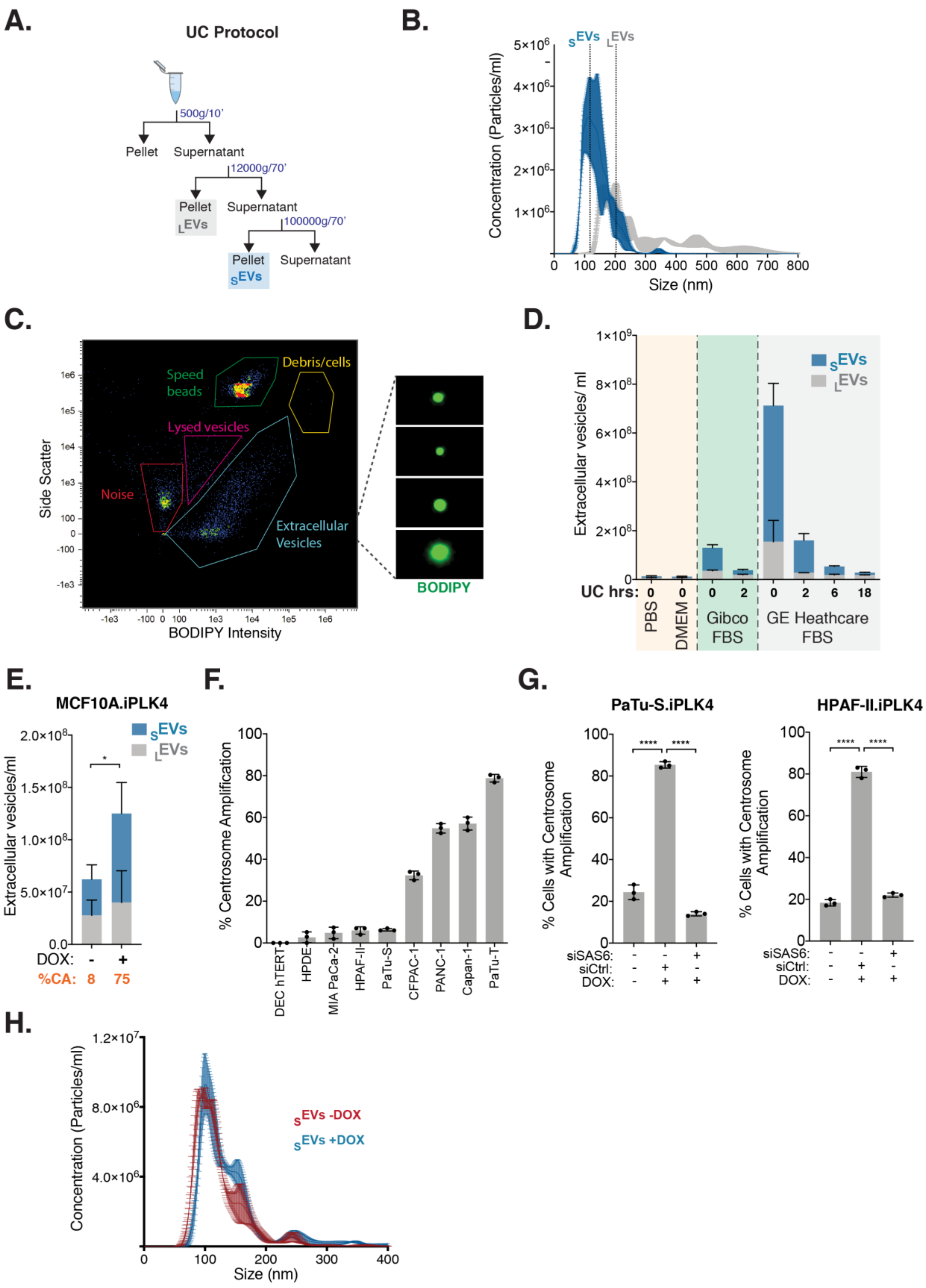


**Figure S1. EV isolation and characterization in PDAC cell lines, related to Figure 1.** (A) Experimental flowchart. (B) Quantification of $_{S}$EVs and $_{L}$EVs concentration and size using the nanoparticle tracking device NanoSight to assess the reliability of the UC protocol to separate EVs by size. (C) Example scatterplot from ImageStream displaying side scatter plotted against BODIPY maleimide intensity. Representative gating regions are shown. Gating region for contaminating cells and cell debris (yellow), speed beads (green) used to internally calibrate the ImageStream and for lysed vesicles (purple) are shown. Representative images or particles taken from the ImageStream image gallery that are present in the EV gating region show spherical BOPIDY-labelled vesicles. (D) Quantification of $_{S}$EVs and $_{L}$EVs in the reagents used to culture cells and prepare EVs for analyses and after UC to determine removal of any contaminant EV. (E) Quantification of secreted $_{S}$EVs and $_{L}$EVs from MCF10A.iPLK4 cell line upon induction of centrosome amplification (+DOX). Average of the percentage of centrosome amplification (CA) per cell line is highlighted in orange. (F) Quantification of the percentage of centrosome amplification in a panel of PDAC cell lines. n=300 mitotic cells for each cell line. (G) Quantification of centrosome amplification in the Patu-S.iPLK4 (left) and HPAF-II.iPLK4 (right) cell lines upon induction of centrosome amplification (+DOX) and Sas-6 depletion by siRNA. n=300 mitotic cells for each condition. (H) Quantification of the size of $_{S}$EVs secreted by Patu-S.iPLK4 cells with (+DOX) and without (-DOX) extra centrosomes using the NanoSight. For all graphics error bars represent mean +/- SD from three independent experiments. *$p < 0.05$, ****$p < 0.0001$. The following statistic were applied: for graph in E two-way ANOVA with Tukey's post hoc test was applied and for graphs in G one-way ANOVA with Tukey's post hoc test was applied.

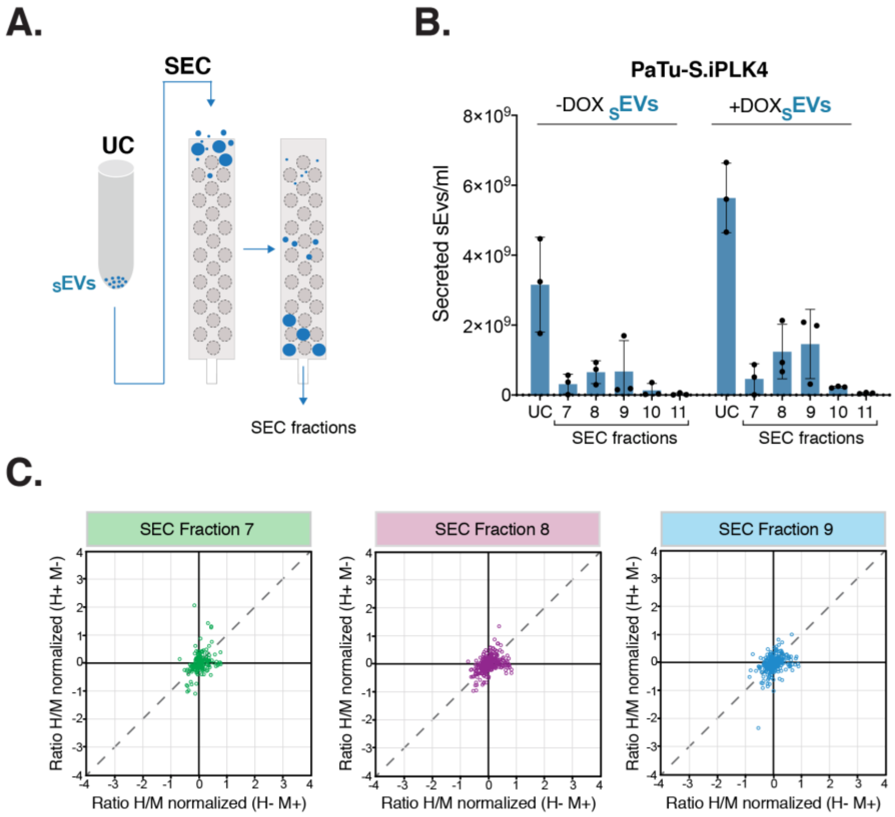


**Figure S2. SILAC proteomic analyses of secreted $_{S}$EVs, related to Figure 2.** (A) Experimental flowchart. (B) Quantification of number $_{S}$EVs in UC and SEC fractions collected from Patu-S.iPLK4 cells with (+DOX) and without (-DOX) extra centrosomes. (C) Correlation graphs plotting $Log_2$ fold change of the ratio of heavy (H) and medium (M) labelled proteins of the forward and reverse experiments for the SEC fractions 7, 8 and 9. Dashed diagonal line illustrates where identical M and H would lie, demonstrating the similarity between H and M labelled $_{S}$EVs. See also Table S4.

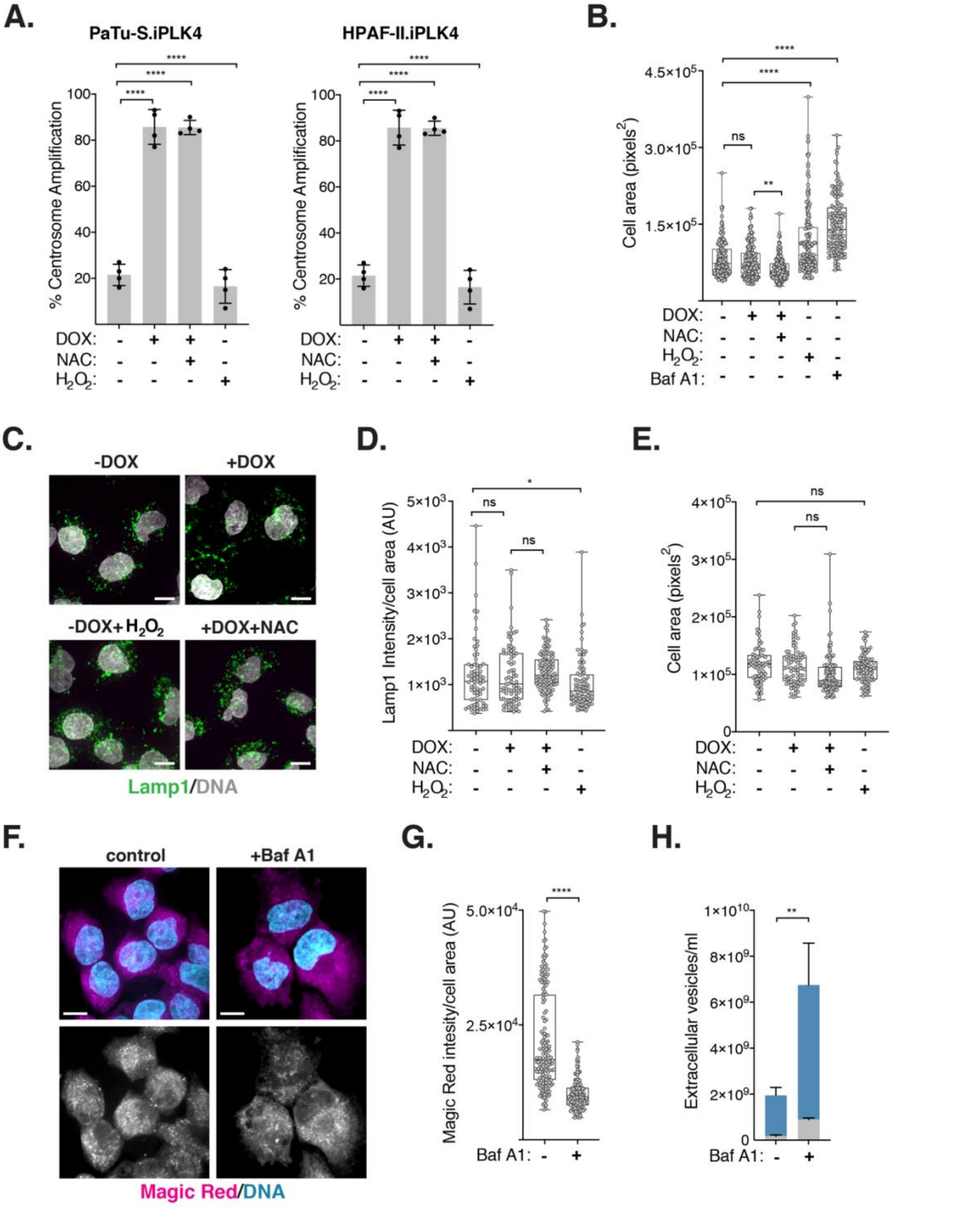


**Figure S3. Characterization of lysosome function in cells with amplified centrosomes, related Figure 3.** (A) Quantification of centrosome amplification in the Patu-S.iPLK4 (left) and HPAF-II.iPLK4 (right) cell lines upon induction of centrosome amplification (+DOX). 5 mM of NAC, 100 µM $H_2O_2$ and 20 nM Baf A1 was used. n=300 mitotic cell lines for each condition. (B) Quantification of cell area (pixels$^2$). 5 mM of NAC, 100 µM $H_2O_2$ and 20 nM Baf A1 was used. $n_{(-DOX)}$=158, $n_{(+DOX)}$=189, $n_{(+DOX+NAC)}$=221, $n_{(-DOX+H2O2)}$=175 and $n_{(+BafA1)}$=144. (C) Representative confocal images of Patu-S.iPLK4 cells stained for total lysosomes (Lamp1, green) and DNA (grey). Scale bar, 10 µm. (D) Quantification of Lamp1 fluorescence intensity in Patu-S.iPLK4 cells normalized for cell area. 5 mM of NAC, 100 µM $H_2O_2$ was used. AU, arbitrary units. $n_{(-DOX)}$=71, $n_{(+DOX)}$=80, $n_{(+DOX+NAC)}$=112 and $n_{(-DOX+H2O2)}$=87. (E) Quantification of cell area (pixels$^2$). 5 mM of NAC, 100 µM $H_2O_2$ was used. $n_{(-DOX)}$=71, $n_{(+DOX)}$=80, $n_{(+DOX+NAC)}$=112 and $n_{(-DOX+H2O2)}$=87. (F) Representative confocal images of Patu-S.iPLK4 cells stained for functional lysosomes (Magic red, magenta) and DNA (cyan) in control cells (-DOX) and cells treated with 20 nM Baf A1. Scale bar, 10 µm. (G) Quantification of intracellular Magic red fluorescence intensity normalized for cell area in Patu-S.iPLK4 cells. AU, arbitrary units. $n_{(control)}$=158 and

$n_{(+BafA1)}$=144. Note that data plotted for control cells is the same as in Figure 3D. (H) Quantification of $_{S}$EVs and $_{L}$EVs secretion in control and Baf A1 treated Partu-S.iPLK4 cells. For all graphics error bars represent mean +/- SD from three independent experiments. *$p < 0.05$, **$p < 0.01$, ****$p < 0.0001$, n.s. = not significant ($p > 0.05$). The following statistic were applied: for graphs in A, one-way ANOVA with Tukey's post hoc test was applied, for graphs in B, D, E and G data one-way ANOVA with a Kruskal-Wallis post hoc test was applied and for data in H two-way ANOVA with Tukey's post hoc test was applied.

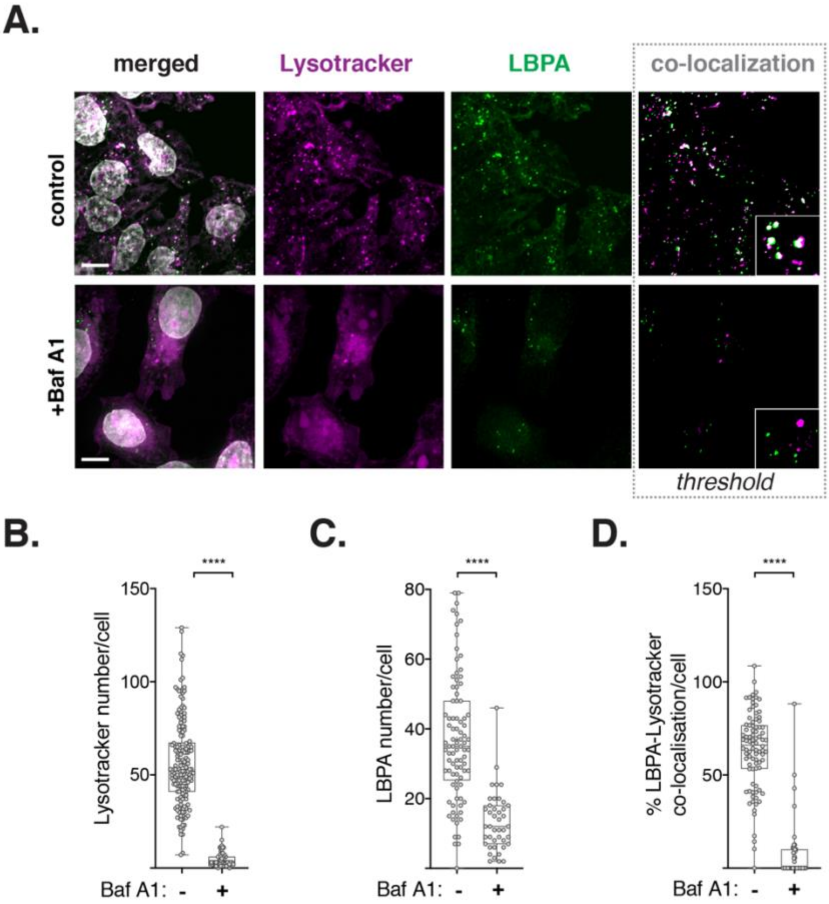


**Figure S4. Bafilomycin A1 treatment disrupts lysosome function and prevents lysosome-MVBs co-localization, related Figure 4.** (A) Representative confocal images of cells stained for acidic lysosomes (Lysotracker, magenta), late endosomes/MVBs (anti-LBPA, green) and DNA (grey). Insets show higher magnification of lysotracker and LBPA-labelled vesicles. Scale bar, 10 $\mu$m. (B) Quantification of the number of lysotracker-labelled lysosomes per cell. 20 nM of Baf A1 was used. $n_{(control)}$=166 and $n_{(+BafA1)}$=67. Note that data plotted for control cells is the same as in Figure 4B. (C) Quantification of LBPA-labelled late endosomes/MVBs per cell. 20 nM of Baf A1 was used. $n_{(control)}$=88 and $n_{(+BafA1)}$=42. Note that data plotted for control cells is the same as in Figure 4C. (D) Quantification of the percentage of lysotracker and LBPA-labelled intracellular vesicles co-localization. 20 nM of Baf A1 was used. $n_{(control)}$=86 and $n_{(+BafA1)}$=42. Note that data plotted for control cells is the same as in Figure 4D. For all graphics error bars represent mean +/- SD from three independent experiments. *$****p < 0.0001$*. Graphs were analyzed with one-way ANOVA with a Kruskal-Wallis post hoc test.

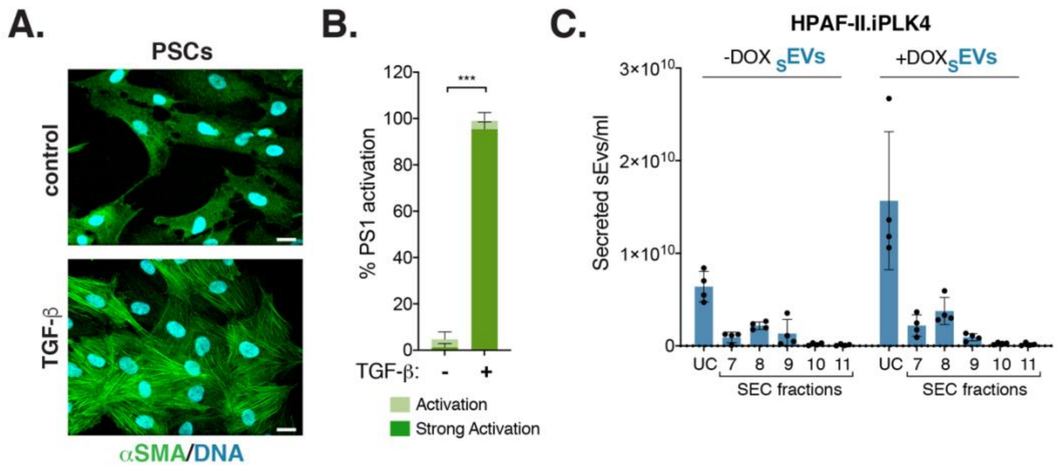


**Figure S5. Characterization of PSCs activation, related Figure 5.** (A) Representative confocal images of PSCs stained for α-SMA (green) and DNA (cyan). Scale bar, 20 μm. (B) Quantification of the percentage of PSCs activation upon treatment with TGF-β, used as positive control. 5 ng/ml of TGF-β was used. PSCs $n_{(control)}$=475, $n_{(+TGF-\beta)}$=414 (C) Quantification of number $_{S}$EVs in UC and SEC fractions collected from HPAF-II.iPLK4 cells with (+DOX) and without (-DOX) extra centrosomes. For all graphics error bars represent mean +/- SD from three independent experiments. *$***p < 0.001$*. Paired *t* test was used for statistical analyses.
